# The COVID-19 mRNA vaccine Comirnaty induces anaphylactic shock in an anti-PEG hyperimmune large animal model: Role of complement activation in cardiovascular, hematological and inflammatory mediator changes

**DOI:** 10.1101/2023.05.19.541479

**Authors:** Bálint András Barta, Tamás Radovits, Attila Balázs Dobos, Gergely Tibor Kozma, Tamás Mészáros, Petra Berényi, Réka Facskó, Tamas Gyula Fülöp, Béla Merkely, János Szebeni

## Abstract

**Background:** Comirnaty, Pfizer-BioNTech’s polyethylene-glycol (PEG)-containing Covid-19 vaccine, can cause hypersensitivity reactions (HSRs) in a small fraction of immunized people which can, very rarely, culminate in life-threatening anaphylaxis. A role of anti-PEG antibodies (Abs) has been proposed, but causality has not yet been proven in an animal model. This study aimed to provide such evidence using anti-PEG hyperimmune pigs (i.e., pigs displaying very high levels of anti-PEG Abs). We also sought to find evidence for the role of complement (C) activation and thromboxane A2 (TXA2) release in blood as contributing effects to anaphylaxis.

**Methods:** Pigs (n=6) were immunized with 0.1 mg/kg PEGylated liposome (Doxebo) i.v. the rise of anti-PEG IgG and IgM was measured in serial blood samples with ELISA. After 2-3 weeks, during the height of seroconversion, the animals were injected i.v. with 1/3 human vaccine dose (HVD) of Comirnaty, and the hemodynamic (PAP, SAP), cardiopulmonary (HR, EtCO2,), hematological parameters (WBC, granulocyte, lymphocyte, and platelet counts) and blood immune mediators (anti-PEG IgM and IgG Abs, C3a and TXA2) were measured as endpoints of HSRs.

**Results:** A week after immunization of 6 pigs with Doxebo, the level of anti-PEG IgM and IgG rose 5-10-thousands-fold in all animals, and they all developed anaphylactic shock to i.v. injection of 1/3 HVD of Comirnaty. The reaction, starting within 1 min, led to the abrupt decline of SAP along with maximal pulmonary hypertension, decreased pulse pressure amplitude, tachycardia, granulo- and thrombocytopenia, and paralleling rises of plasma C3a and TXB2 levels. These vaccine effects were not observed in non-immunized pigs.

**Conclusions:** Consistent with previous studies with PEGylated nano-liposomes, these data show a causal role of anti-PEG Abs in the anaphylaxis to Comirnaty. The reaction involves C activation, and, hence, it represents C activation-related pseudo-allergy (CARPA). The setup provides the first large-animal model for mRNA-vaccine-induced anaphylaxis in humans.

## INTRODUCTION

Anaphylaxis caused by the mRNA-lipid nanoparticle (mRNA-LNP)-based Covid-19 vaccines, Comirnaty and Spikevax, in the range of 3-234 cases per million vaccinees, is considered as a rare adverse event.^1-17^. Regarding its mechanism, allergy to PEG is one reason,^18^ however, there is consensus in the literature that the overwhelming majority of cases are not representing classical type-1 allergy but are IgE-independent “pseudoallergies”.^9, 10, 16, 19^ These allergy-like reactions arise without prior sensitization, by way of direct and/or indirect stimulation of mast cells and also, macrophages, platelets, granulocytes, which are typically not involved in Type-1 allergy.^20-23^ The latter cells can be activated both by direct binding of reactive NPs to their inflammatory signal receptors, and/or anaphylatoxins (C3a, C5a) to their specific surface receptors (C3aR, C5aR). Because anaphylatoxins are byproducts of complement (C) activation, the reactions proceeding through the latter pathway are referred to as C activation-related pseudoallergy (CARPA)^24^ although C activation may be only a co-trigger.^25^

In support of the involvement of CARPA in Comirnaty-induced HSRs we and others have pointed out that multiple components of the vaccine can activate C via different pathways.^26,27^ Notably, nucleic acids can trigger C activation via the classical pathway; cationic lipids via the alternative pathway and the spike protein, via the lectin pathway.^26 27^ In addition, the PEGylated LNP carrier can also trigger classical pathway C activation due to the binding of anti-PEG Abs to the PEG on the NP surfaces, for which astonishing visual and functional evidence was served recently by demonstrating the membrane damage caused by the C membrane attack complex (C5b-9).^29, 30^ Along this line, Comirnaty turned out to be a strong activator of porcine C,^28^ and i.v. administration of the vaccine in pigs mimicked the typical hemodynamic, hematological and blood TXB2 changes seen in liposome-induced CARPA in pigs.^28^

Regardless of C activation, the likely causal role of anti-PEG Abs in mRNA-LNP vaccine-induced HSRs/anaphylaxis was raised by many groups,^1, 8, 15, 16, 18, 31^ and we found significant correlation between the blood levels of anti-PEG Abs and rise of HSR/anaphylaxis in recipients of Comirnaty and Spikevax.^32^ Nevertheless, all above facts provide only indirect proofs of a causal role of anti-PEG Abs in Comirnaty-induced HSRs; conclusive experimental or clinical evidence is still lacking. Accordingly, the goal of the present study was to obtain such evidence using the anti-PEG hyperimmune pig model,^33^ which showed the acceleration of HSR to anaphylaxis to PEGylated liposomal doxorubicin (Doxil) if the blood level of anti-PEG IgM had been increased by prior vaccination with drug-free Doxil (Doxebo).^33^

## METHODS

### Materials

Comirnaty was from Pfizer/BioNtech, the vaccine used for human vaccinations against SARS-Cov-2 infections. One 0.3 mL shot contains, in addition to phosphate buffer and sucrose, 30 μg mRNA, 430 μg ALC-0315, (4-hydroxybutyl)azanediyl)bis(hexane-6,1-diyl)bis(2-hexyldecanoate)}; 50 μg ALC-0159, 2-[(polyethylene glycol)-2000]-N,N ditetradecylacetamide; 90 μg 1,2-Distearoyl-sn-glycero-3-phosphocholine (DSPC) and 200 μg cholesterol. Its total lipid content is: 0.77 mg. The porcine C3a kit was obtained from TECO*Medical* AG, Sissach, Switzerland (Cat No: TE1078) and the TXB2 ELISA was from Cayman Chemical (Ann Arbor, MI, USA). Zymosan, Dulbecco’s phosphate-buffered saline (PBS) without Ca2+/Mg2+ and bovine calf serum, and biotin-labeled goat polyclonal anti-porcine IgM were from Sigma Chemical Co. (St. Louis, MO, USA).

### Preparation of Doxebo

The preparation and characteristics of Doxebo were described earlier in detail.^33^ In brief, the freeze-dried lipid components of Doxil were hydrated in 10 mL sterile pyrogen-free normal saline by vortexing for 2−3 min at 70°C to form multilamellar vesicles (MLVs). The MLVs were downsized through 0.4 and 0.1 μm polycarbonate filters in two steps, 10 times through each, using a 10 mL extruder barrel from Northern Lipids (Vancouver, British Columbia, Canada) at 62 °C. Liposomes were suspended in 0.15 M NaCl/10 mM histidine buffer (pH 6.5). The size distribution (Z-average: 81.17 nm) and phospholipid concentration of Doxebo (12.6 mg/mL) were determined as described earlier.^33^

### Animals

Mixed breed Yorkshire/Hungarian White Landrace pigs of both sexes (2−3 months old, 20−28 kg) were obtained from the Animal Breeding and Nutrition and Meat Science Research Institute, Hungarian University of Agriculture and Life Sciences (Herceghalom, Hungary).

### Treatment protocol

As outlined in Fig 1, baseline (“pre-Immun”) blood samples were taken from 6 pigs followed by immunization by way of infusion of 0.1 mg PL/kg Doxebo via the ear vein (suspended in 20 mL of saline) at a speed of 1 mL/min. The animals were then placed back into their cages until the 2nd blood sampling 9-10 days later, to screen for anti-PEG Ab induction. From 3 days later, within a period of 13 days, the animals showing seroconversion (all 6 of 6) were subjected to the “CARPA induction” protocol with Comirnaty as well as one control animal which was sham-immunized with PBS but was handled identically to the Doxebo-immunized pigs.

**Figure 1.**
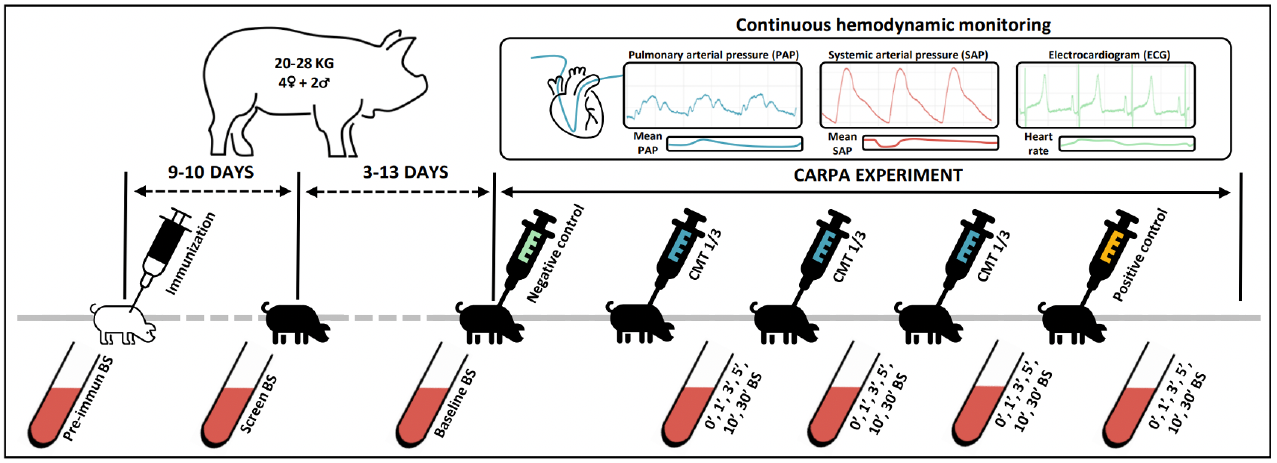
Timeline of the experimental protocol testing the physiological effects of i.v. Comirnaty in anti-PEG hyperimmune pigs.

In order to induce CARPA, the animals were sedated with a mixture of ketamine and xylazine (intramuscularly), then anesthesia was induced with isoflurane (2−3% in oxygen in 100% FiO2). This was followed by intubation with endotracheal tubes (ET tube size: 6-7, ET cuff pressure: <25 cmH2O) and inhalation anesthesia was maintained with isoflurane (1.5−2% in oxygen 100% FiO2). In addition, local anesthesia was performed at the site of interventions with lidocaine. In case apnea occurred during the experiment, assisted mechanical ventilation was initiated (in volume mode, Vt.: 220-320 ml, Rate: 16-18/m). After surgical scrubbing of the skin, with detergent and disinfectants (BradoNett disinf. liquid soap and 10% povidone-iodine (Betadine sol.)), the pigs were subjected to surgery to insert various catheters into their circulation, namely: (a) a Swan−Ganz catheter (Arrow AI-07124 single-lumen balloon wedge pressure catheter 5 Fr., 110 cm, Arrow International Inc, Reading, PA, USA), into the pulmonary artery via the right external jugular vein (in order to measure the pulmonary arterial pressure (PAP); (b) the left femoral artery to record the systemic arterial pressure (SAP); (c) the left external jugular vein for saline and drug infusion; (d) into the left femoral vein for blood sampling; and (e) the right common carotid artery for arterial blood gas analysis. After 15-30 min adaptation, the 6 Doxebo-immunized animals were treated by 5 consecutive i.v. injections into left external jugular vein of (1) 5 mL saline (to provide baseline for the hemodynamic changes), (2) bolus injection of 1/3 HVD of Comirnaty (to trigger the immune reaction), (3, 4) 2 repeats of the same dose of Comirnaty (to establish tachyphylaxis, i.e., self-induced tolerance), and finally (5) with 0.1 mg/kg zymosan, as positive control. One control sham-immunized animal underwent steps (1) (saline, negative control), (2) bolus injection of 5X human dose of Comirnaty and (5) (0.1 mg/kg zymosan, positive control).

All injections were administered under 30 seconds. Among the injections 15-60 min breaks were taken to allow the hemodynamic parameters return to baseline. The latter, as well as the ECG data, were recorded by the physiological monitoring systems of Pulsion Medical Systems SE (Munich, Germany) and Powerlab (ADInstruments, Bella Vista, Australia). The arterial blood gas analysis was executed with a Roche COBAS B221 benchtop analyzer (Roche Diagnostics, Rotkreuz ZG, Switzerland). End-tidal pCO2, O2 saturation, ventilation rate and body temperature were also continuously measured. At the end of the experiments the animals were sacrificed with pentobarbital (120mg/kg iv.) and concentrated potassium chloride.

### Blood cell assays

For the blood cells assays 10 mL venous blood samples were drawn from the pigs at different times into EDTA containing vacuum blood collection tubes (K3EDTA Vacuette, Greiner 367 Bio-One Hungary, Mosonmagyaróvár, Hungary) and aliquoted to 0.5 mL Eppendorf tubes. The white blood cell (WBC), granulocyte (GR), lymphocyte (LY), platelet (PLT) and red blood cell (RBC) counts and hemoglobin (Hgb) concentration were determined using an ABACUS Junior Vet hematology analyzer (Diatron, Budapest, Hungary).

### ELISA of Anti-PEG antibodies

For the analysis of anti-PEG Abs, blood was taken from the ear vein before pretreatment and then at different times specified in the Results. Anticoagulation was done with K3-EDTA tubes. Polysorp (Nunc) plates were coated with 1.25 μg/well DSPE-PEG-2000 in 100 μL of bicarbonate buffer (4.46 μM) (pH ∼9.0) overnight at 4 °C, followed by blocking of the wells with 150 μL of PBS/0.05% Tween-20 + 2% bovine serum albumin (BSA) at 37 °C for 1.5 h. Before and after blocking, wells were washed two and three times with 300 μL of wash buffer containing PBS/0.05% Tween-20 for 1 min, respectively. The EDTA-anti-coagulated plasma samples were diluted by PBS/0.05% Tween-20 + 1% BSA in the 10−19 500-fold range and incubated in the wells for 1.5 h at 37 °C, with slow shaking. Wells were washed five times with 300 μL of wash buffer for 1 min. After staining with 100 μL of HRP-conjugated anti-porcine IgM (2000× dilution, Sigma) or IgG (800× dilution, Sigma) for 1 h, wells were washed again five times with wash buffer as mentioned. The antibodies were stained by incubation with 100μL of substrate solution (Neogen) containing 3,3′,5,5′-tetramethylbenzidine (TMB) and hydrogen peroxide for 15 min in dark. The reaction was stopped with 50 μL of 2 N H2SO4, and A450 was read with a Fluostar Omega 96-well plate reader (BMG Labtech, Ortenberg, Germany). The titer unit was defined as the dilution at which the blank-corrected OD was 0.1.^5^

### ELISA of blood levels of TXB2 and C3a

To measure thromboxane B2 (TXB2), a stable metabolite of thromboxane A2 (TXA2), 4 μg indomethacin (diluted in 2 ul of 96% ethanol) was mixed to 2 mL of EDTA-anticoagulated blood to prevent TXA2 release from WBC before centrifugation at 2000g, for 4 min at 4°C. The plasma samples were aliquoted, frozen, and stored at -70°C until the TXB2 assay was performed using a kit from Cayman Chemicals (Ann Arbor, MI, USA) and a FLUOstar Omega microplate reader (BMG 379 Labtech). Plasma levels of porcine C3a was measured in EDTA-anticoagulated blood samples, using a porcine specific C3a ELISA kit obtained from TECO*Medical* AG, Sissach, Switzerland (Cat No: TE1078).

### Statistical Methods

Values at all time points were compared to their baseline (−1 min), and the significance of differences was determined by nonparametric Paired Samples Wilcoxon test. Depletion of immunoglobulins was tested with Trend analysis carried out in Graphpad Prism. Reactions to repeated injections of 1/3 HVD of Comirnaty were compared with Friedman-test, followed by Wilcoxon post-hoc test. Correlation among parameters was examined with Spearman’s method. A P-value of <0.05 was considered statistically significant.

### Ethics

The investigation conformed to the EU Directive 2010/63/EU and the Guide for the Care and Use of Laboratory Animals used by the US National Institutes of Health (NIH Publication No.85– 23, revised 1996). The experiments were approved by the Ethical Committee of Hungary for Animal Experimentation (permission number: PE/EA/843-7/2020).

## RESULTS

### Raising of blood anti-PEG Ab levels by immunization with Doxebo

Fig 2A and C show the absolute levels of anti-PEG IgM and IgG on days 0 (Pre-Immune) and 9-10 (Screen) on a logarithmic scale. Immunization was successful in all animals, although with some individual variation. Fig 2B, D displays IgM and IgG levels on the day of the experiment 12-23 days after immunization with Doxebo, in blood samples taken before the first (r0), second (r1) and third (r2) injections of 1/3 HVD of Comirnaty.

**Figure 2.**
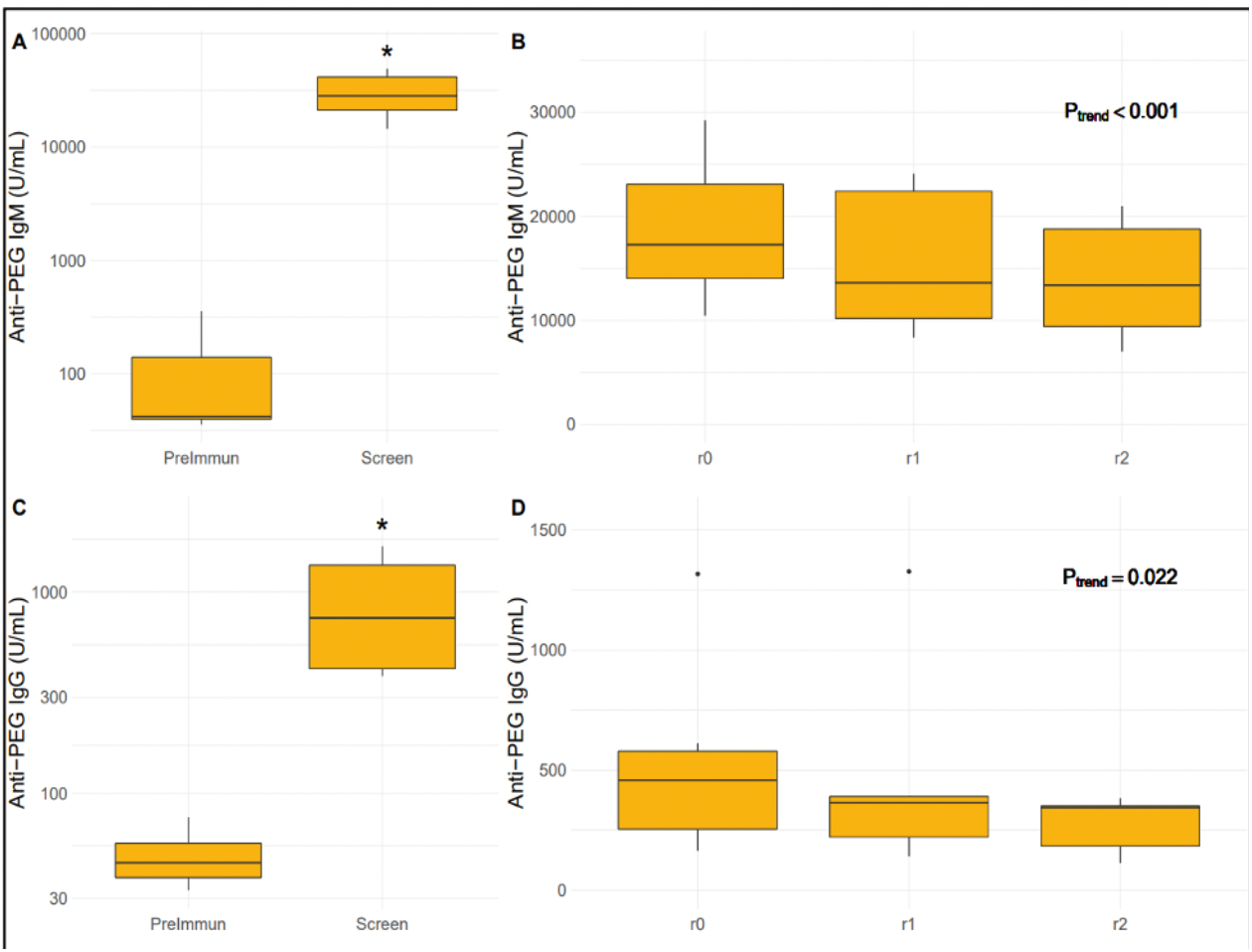
Panels A and C shows the absolute levels of anti-PEG IgM and IgG on a logarithmic scale just before (PreImmun) and 9-10 days after (Screen) immunization with Doxebo. Panels B and D show anti-PEG IgM and IgG levels on the day of the experiment before the first (r0), second (r1) and third (r2) injection of 1/3 HVD of Comirnaty.

These data demonstrate that the immunization was effective in each animal, implying that the Comirnaty challenge on the 12-23 days postvaccination interval was performed in anti-PEG Ab hyperimmune animals. Furthermore, our analysis has identified a significant downward trend in the blood levels of both Abs, suggesting their progressive depletion as a consequence of repeated exposure to Comirnaty.

### Induction of anaphylaxis by Comirnaty in anti-PEG hyperimmune pigs: Characteristics of the reaction

Figure 3 shows that 1/3 HVD of Comirnaty caused robust hemodynamic changes leading to shock within 1-2 min after i.v. injection. The reaction involved maximal rise of PAP within 1 min after the vaccine’s injection, with initially unchanged pulmonary arterial pulse pressure amplitude (A-B), which was paralleled by an abrupt decline in systemic arterial pulse pressure amplitude shortly followed by falling SAP (C-D). The 3-second snapshots from the third minute after the injection highlight the massive signal distortions of PAP, SAP and ECG wave morphology due to cardiopulmonary resuscitation (CPR) involving chest compressions and noradrenaline administration followed by tachycardia and rebound systemic hypertension (B, D, F).

**Figure 3.**
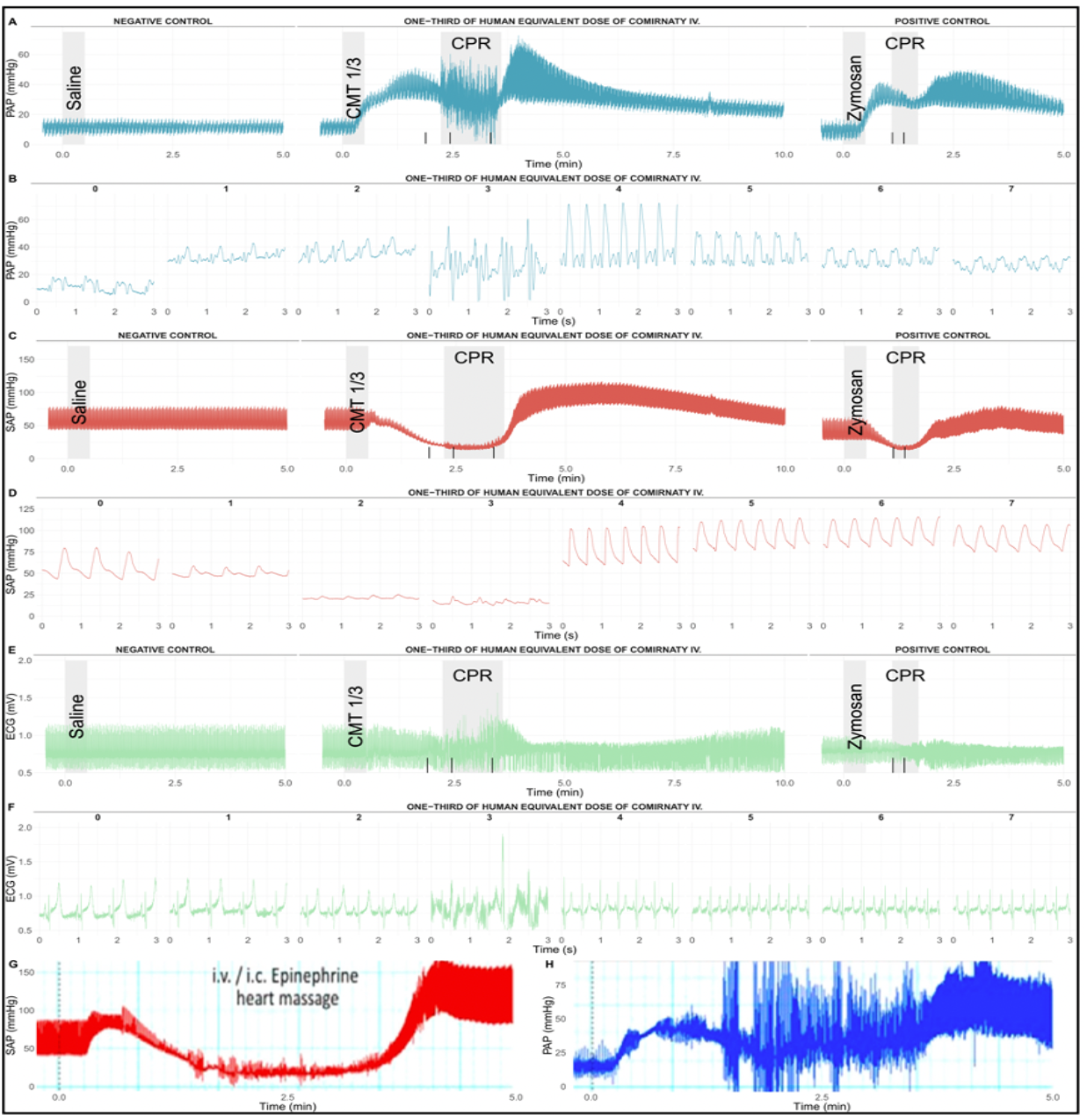
Typical real-time hemodynamic and ECG tracings in 1 of 6 pigs injected i.v. with 1/3 HVD of Comirnaty 16 days after immunization with 0.1 mg/kg Doxebo, as described in the Methods. Saline (PBS) bolus to establish the baseline; vertical black lines correspond to injections of noradrenaline in 1:100 dilution during CPR. CMT 1/3, i.v. bolus injection of 1/3 human dose of Comirnaty; CPR, cardiopulmonary resuscitation; Zymosan, bolus injection of 0.1 mg/kg zymosan.

The CPR salvaged the animal, which could be later injected two more times (2 repeat vaccine injections, see below), and finally with 0.1 mg/kg zymosan. The latter caused similar, although less intense hemodynamic changes, which implies that the vaccine’s effect was similar or stronger than that of the standard positive control zymosan at a 83-fold lower dose in terms of weight. Fig 3G-H reproduces the SAP and PAP changes caused by Doxebo in anti-PEG hyperimmune pigs,^33^ highlighting the practical identity of vaccine- and liposome-induced reactions that was considered as pseudo-anaphylaxis.^33^

Figure 4 summarizes the hemodynamic alterations in 6 pigs after the first injection of 1/3 HVD of Comirnaty. Boxplots of mean pressure and pulse pressure derived from PAP and SAP (A-D), as well as HR, R wave amplitude and ST height derived from the ECG signal (E-G) are shown. The highly reproducible, statistically significant rises of mean PAP and HR, as well as the declines of mean SAP, R amplitude and ST heights, are typical symptoms of CARPA in this model.^24, 34-38^ Panel H highlights the pathologic proximity of mean PAP and SAP 1.5 minutes into the reaction, i.e., near identity of blood pressures in the pulmonary and systemic circulation. Furthermore, the marked shrinkage of pulse amplitude of SAP seems to occur immediately, detected as early as 0.5 minutes after injection of Comirnaty, while a significant drop in mean SAP was only detected at 1.5 minutes. On the contrary, mean PAP rose without delay upon injection of 1/3 HVD of Comirnaty, while the pulse amplitude of PAP was unaffected by the ongoing reaction, surging only after successful CPR. The sluggishness of decline in mean SAP may postpone the detection of a severe reaction, leaving less chance for timely decision of initiation of CPR. Since in a clinical setting, measurement of PAP is problematic, monitoring of early changes in the pulse amplitude of SAP may serve as a better early warning tool.

**Figure 4.**
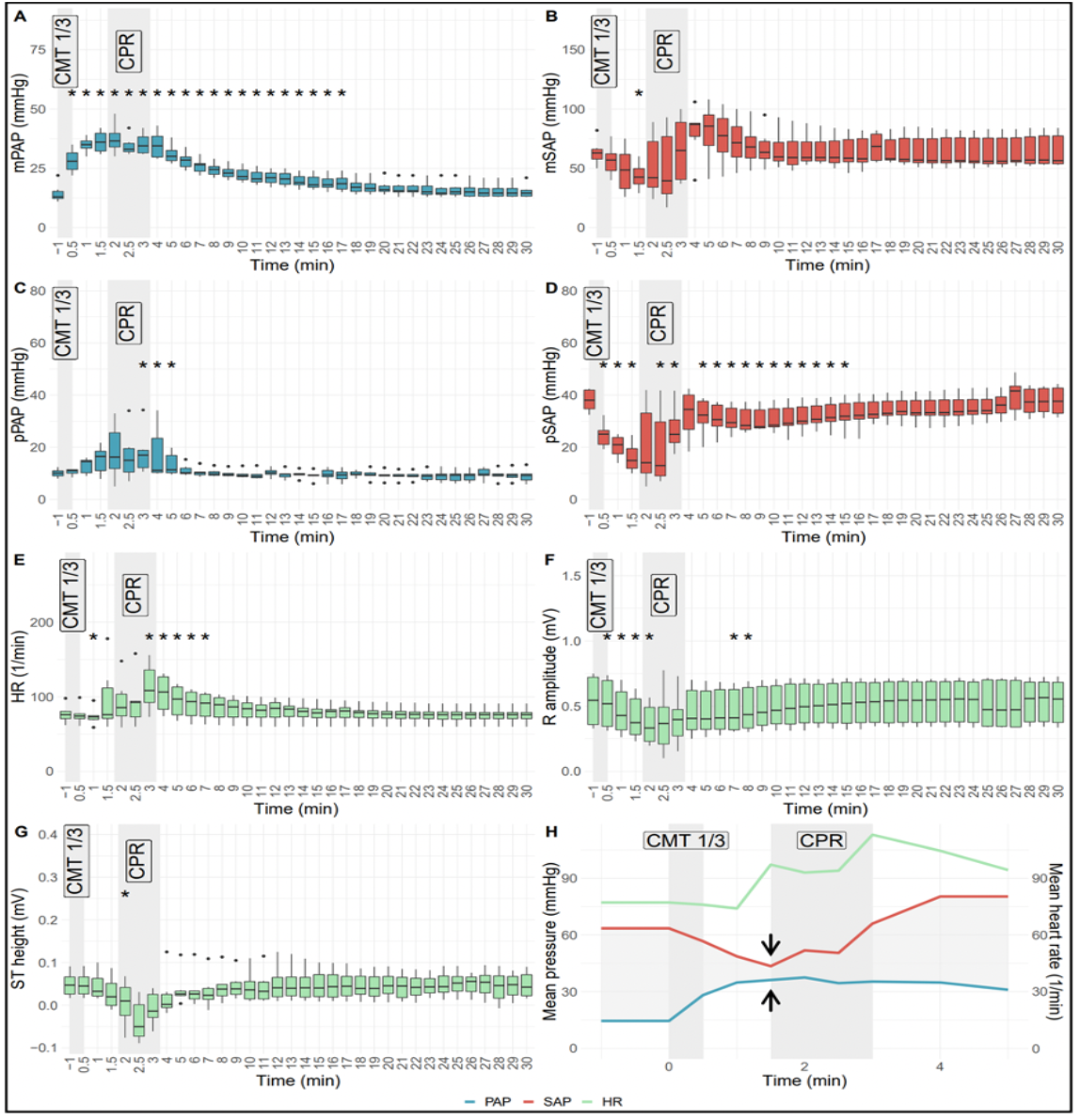
Boxplots of hemodynamic and ECG changes in 6 Doxebo-immunized, anti-PEG hyperimmune pigs injected i.v. with 1/3 HVD of Comirnaty 12 - 23 days after treatment with 0.1 mg/kg Doxebo, as described in the Methods. mPAP, mSAP, pPAP, pSAP denote mean and pulse pressure of PAP and SAP. The opposing arrows on panel H highlight the near equivalence of PAP and SAP 1.5 minutes after the injection. All other abbreviations and labels are the same as in Figure 3. Values at all time points were compared to their baseline (−1 min), and the significance of differences was determined by nonparametric Paired Samples Wilcoxon test (*, p<0.05).

### Comirnaty-induced hemodynamic changes are partly tachyphylactic in anti-PEG hyperimmune pigs

Figure 5 shows the effects of 2nd and 3rd repeat injections of 1/3 HVD of Comirnaty beside the 1st injection, all expressed as % of baseline. The repeated injections entailed significant decrease of PAP and SAP responses whose first signs were diminished increase of mean PAP and reduced decrease of the pulse amplitude of SAP. Thus, tachyphylaxis was initially only partial, and became full only after the second repeat injection.

**Figure 5.**
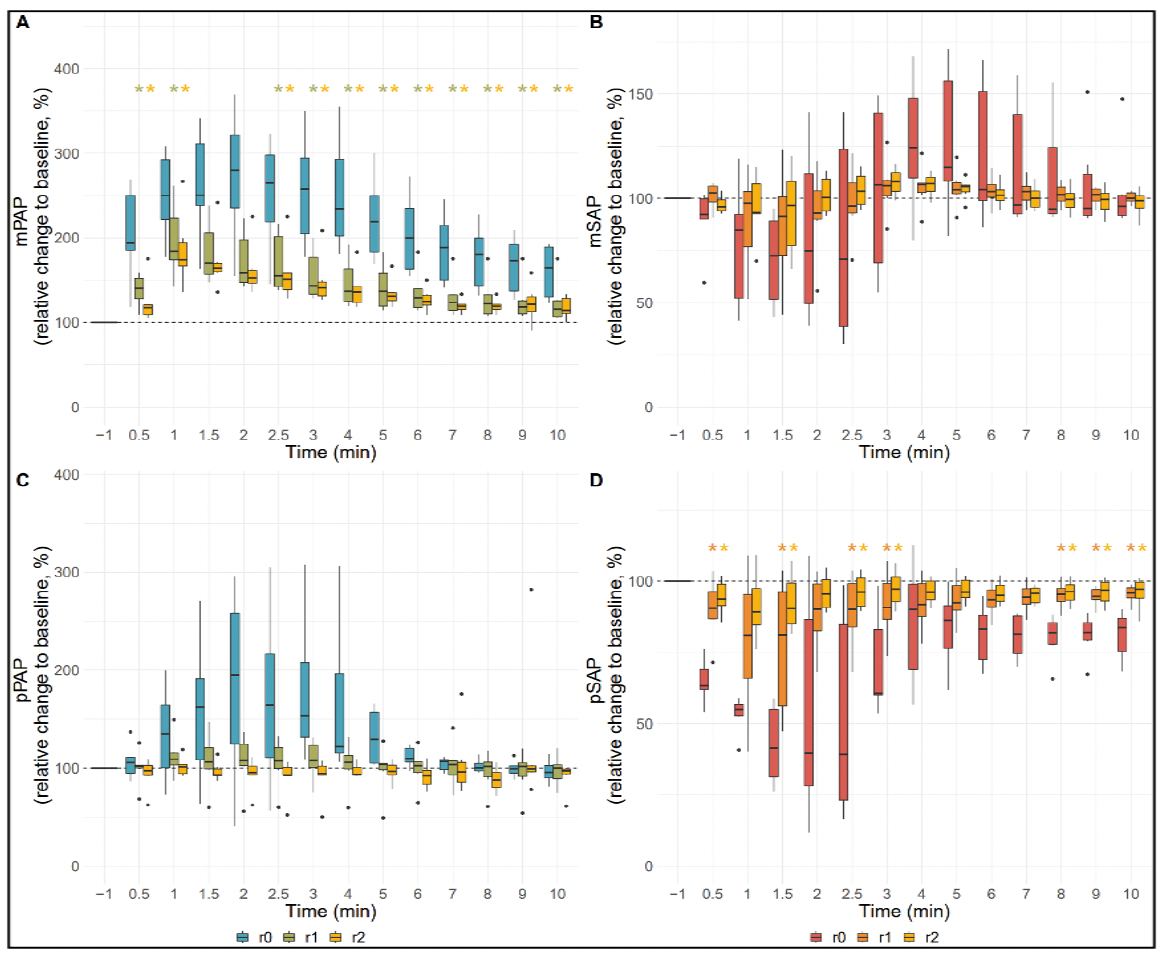
Boxplots showing the gradual decrease of cardiopulmonary response to 1/3 HVD of Comirnaty in 6 anti-PEG hyperimmune pigs. Key: r (repeat) 0, r1 and r2 represents the 1st, 2nd and 3rd injection of the vaccine. Values normalized to baseline preceding each injection (−1) are displayed. Reactions to repeated injections of 1/3 HVD of Comirnaty were compared with Friedman-test, followed by Wilcoxon post-hoc test. *, significantly (p<0.05) ameliorated r1 or r2 response due to partial tachyphylaxis compared to r0. Coloring of the * corresponds to the repeat reaction (r1 / r2) with the same color value.

Figure 6A-B and C-H show the respiratory and hematologic endpoints, respectively. Among these, the significant drops of etCO2 (A), platelet (E) and WBC counts (F) are also typical symptoms of CARPA,^24, 34-38^ while the lack of changes in oxygen saturation (B), RBC count (C), hemoglobin (D) and relative abundance of granulocytes (G) or lymphocytes (H) are not known to be CARPA-dependent variables on the time scale of minutes.

**Figure 6.**
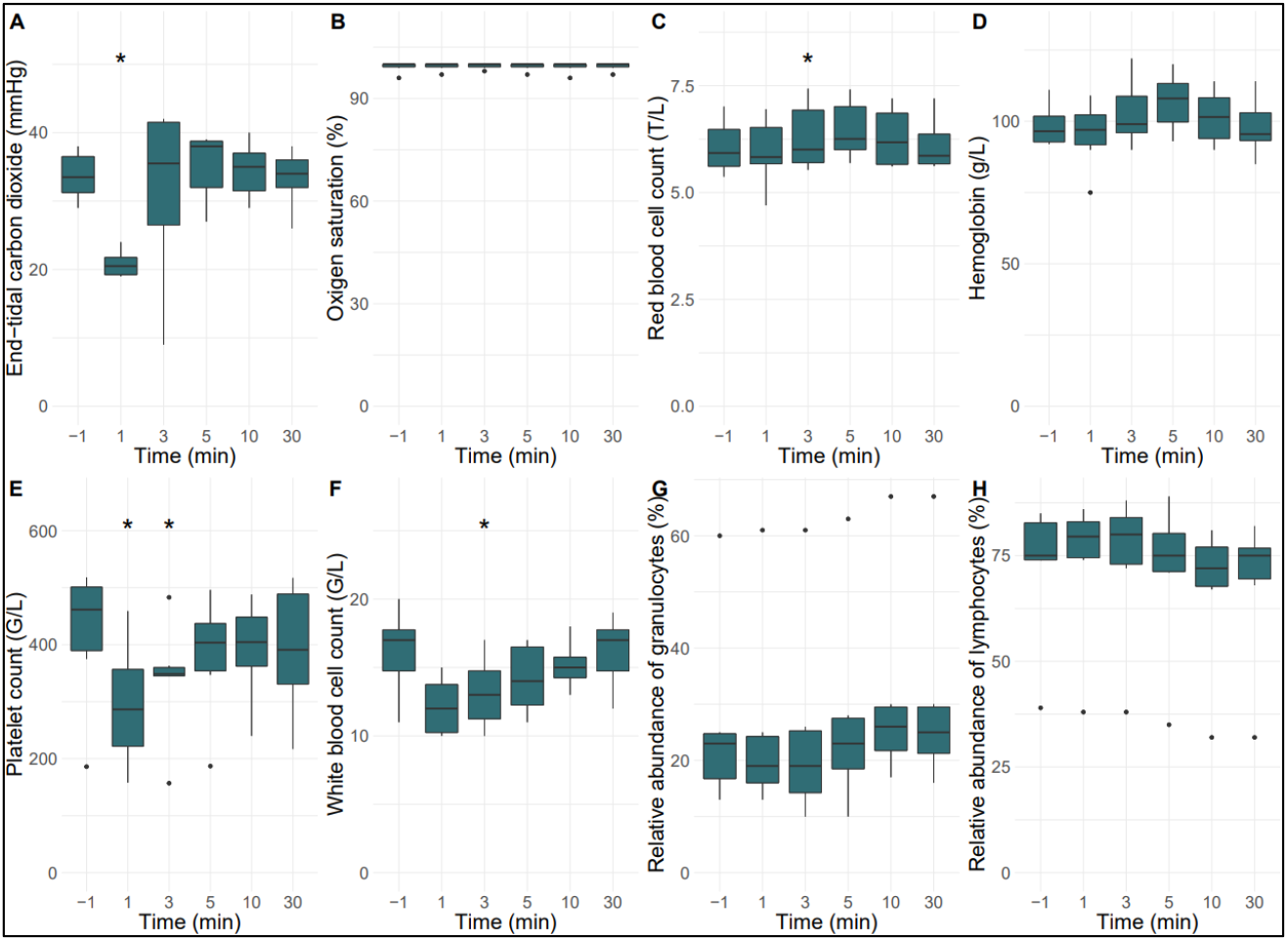
Summarized respiratory (A, B*) and hematologic (C-H) changes in 6 Doxebo-immunized, anti-PEG hyperimmune pigs injected i.v. with 1/3 HVD of Comirnaty. All other details are explained in the Methods and other figure legends. Values at all time points were compared to their baseline (−1 min), and the significance of differences was determined by nonparametric Paired Samples Wilcoxon test (*p<0.05). B*, The ventilation with 2-3% isoflurane in O2 further stabilized the O2 saturation.

The above observations, taken together with the absence of, or minimal physiological changes caused by 5X HVD of Comirnaty in 8 of 9 Doxebo-non-immunized, naive pigs in our previous study,^28^ and the negligible physiological changes in the sham-immunized pig in the present study (Supplementary SFig. 1) provide strong evidence for that very high anti-PEG Ab levels play an essential role in Comirnaty-induced anaphylaxis in pigs.

### Comirnaty-induced changes in plasma C3a and TXB2

To address the question, how anti-PEG Abs could trigger anaphylaxis, we have measured the plasma levels of C3a and TXB2 during the reactions, which inform about the involvement of C activation and the release of TXA2, respectively. Figure 7 shows the time course of changes of plasma TXB2 and C3a after Comirnaty administration in anti-PEG hyperimmune pigs, both inflammatory mediators rising and declining on the same time course of minutes, in close parallelism with the hemodynamic changes. Levels of pSAP and mPAP, the two most sensitive parameters of the CARPA reaction evoked by the injection of 1/3 HVD of Comirnaty showed significant correlation with C3a and TXB2 as well (Fig. 7).

**Figure 7.**
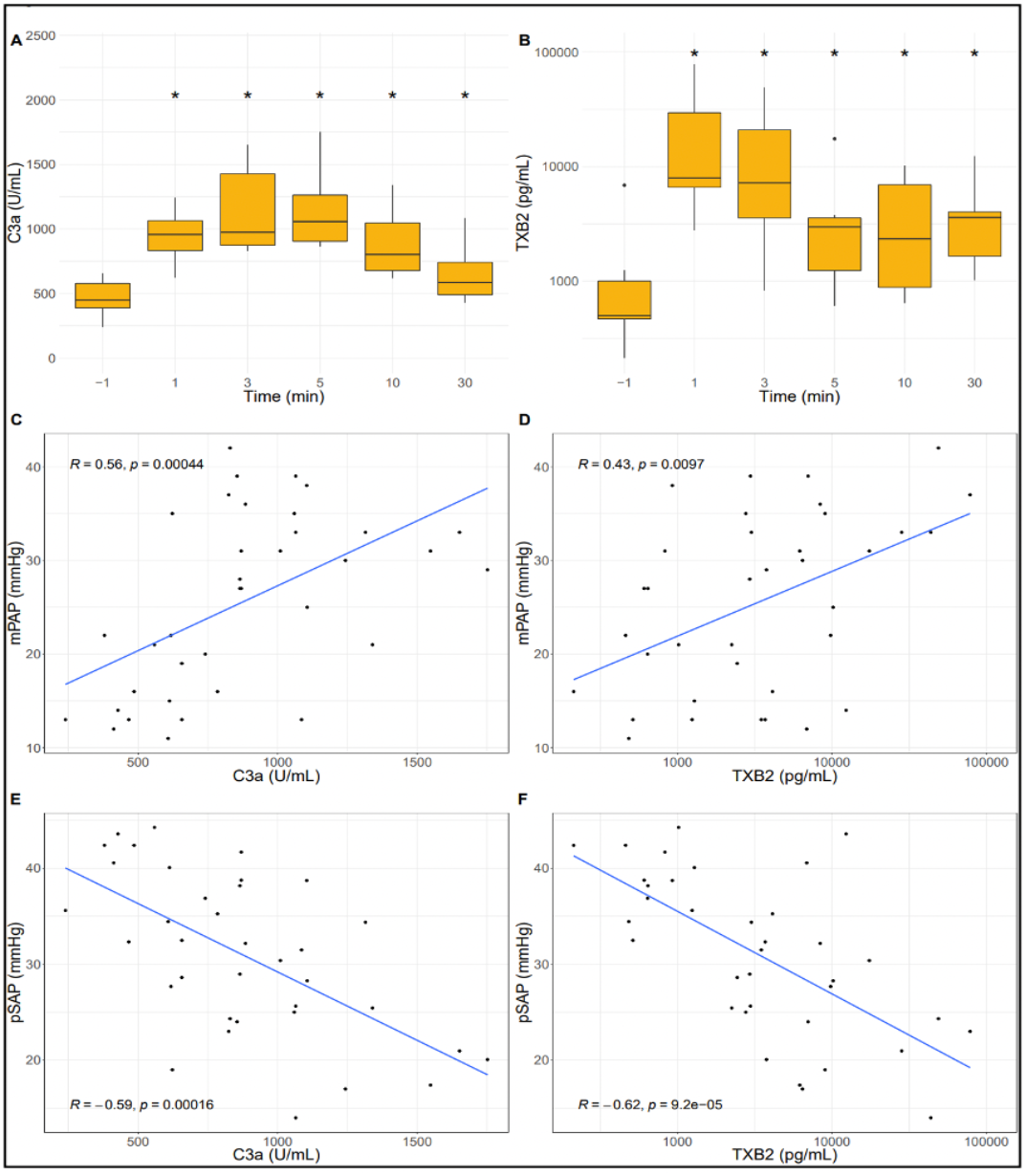
Boxplots of the time course of C3a (A) and TXB2 (B) rise following first i.v. injection of 1/3 HVD of Comirnaty in anti-PEG hyperimmune pigs. TXB2 data is shown on logarithmic scales. The injection of the vaccine resulted in significant elevation of C3a and TXB2 levels. Values at all time points were compared to their baseline (−1 min), and the significance of differences was determined by nonparametric Paired Samples Wilcoxon test (*p<0.05). The figure shows the Spearman correlation of mean pulmonary arterial pressure (mPAP) and systemic arterial pulse pressure (pSAP) with C3a and TXB2 levels.

These data, particularly the identical time course of C3a, TXB2, mPAP and pSAP changes on the minute scale, provide direct evidence for that C activation and subsequent inflammatory cell activation are causally involved in the hemodynamic changes; the reaction proceeds through an immune-cardiovascular axis.

## Discussion

### Clinical relevance

Beyond the clear merits of COVID-19 vaccinations in reducing the morbidity and mortality of SARS-CoV-2 infections, the record number of vaccinations worldwide brought along another scientific benefit, namely, new insights into the mechanism of vaccine efficacy and occasional side effects. This study focused on one adverse effect of the mRNA-LNP vaccine, Comirnaty; an allergic reactivity manifested in HSRs that occasionally culminate in anaphylaxis. The increased risk for anaphylactic reactions to Comirnaty was recognized soon after the introduction of the vaccine in December, 2019,^31^ leading to the exclusion of people with allergy to a vaccine component, or because of genetic proneness for anaphylaxis. Yet, anaphylactic reactions to Comirnaty continued to occur; in fact, they have been listed as number 1 on the manufacturer’s most recent adverse effect list,^39^ and anaphylaxis is only the top of the HSR iceberg.

Regarding the prevalence of anaphylaxis, which entails death or disability in 1.7%,^40^ the ∼1.8 billion mRNA-LNP injections given worldwide in 3 years places even the lowest estimate of the sheer number of anaphylaxis cases in the multiple thousand range. Calculating with the median of estimated anaphylaxis rate (123 cases/million),^32^ yields 223,200 anaphylaxis with ∼3,800 death or disability worldwide, placing vaccine-induced anaphylaxis as the first pandemic of an iatrogenic rare (orphan) disease. To prevent its occurrence in the future, when new infection diseases and vaccines arise, the phenomenon needs to be understood. The present study aimed to pursue this goal.

### The anti-PEG hyperimmune pig CARPA model

Since its first description in 1999,^24^ pigs have been used to study liposome- and other NP-induced HSRs.^28, 33-35, 41-45^ Although criticized for overt sensitivity, this feature is uniquely beneficial when rare diseases need to be studied,^38^ such as the infusion reactions to liposomal drugs, or vaccine-induced HSRs/anaphylaxis. Among these pig studies in the past, two directly led to the present investigation. In the first, the mechanism of doxorubicin-containing PEGylated liposome (Doxil)-induced HSRs was studied, and we immunized pigs with placebo (drug-free) Doxil, named Doxebo, to induce the rise of anti-PEG Abs in blood.^33^ This treatment led to several thousand-fold rise of blood anti-PEG IgM level in 6-7 days, at which time both Doxil and Doxebo caused life-threatening anaphylactic shock in all (5/5) animals within minutes after i.v. administration. The second study^28^ explored the pigs’ response to i.v. administered Comirnaty and found that i.v. administration of 5X HVD of Comirnaty caused typical CARPA symptoms in 3 of 9 animals, with 1 anaphylaxis.^28^ However, these were naive animals regarding anti-PEG immunity, and the blood levels of anti-PEG Abs were low and highly variable, which we could not correlate with the reactions. Fusing on the reactogenicity of Comirnaty in PEG-hyperimmune versus naive animals was expected to provide direct evidence for a causal role of anti-PEG Abs in anaphylaxis, and, as the data showed, it is in fact what happened.

The model applied in the present study has many unique benefits. One is that further sensitization of already sensitive pigs for CARPA provides a model for those people who have very high anti-PEG Ab levels (i.e., anti-PEG Ab supercarriers),^32^ and, hence, are prone not only to vaccine-induced HSR/anaphylaxis but also to those caused by many other PEGylated drugs and household items. Yet another unique benefit is that the hemodynamic and cardiopulmonary changes mimic those human circulatory abnormalities (mainly cardiopulmonary distress, acute myocardial infarction), that make cardiac anaphylaxis life-threatening. Furthermore, the high anti-PEG Ab levels represents functional reproduction of severe pseudoallergy, which increases the risk of anaphylaxis to Comirnaty.^13^

On the contradictory side, the present model deviates from the human vaccination practice in that we administered the vaccine i.v., while people are vaccinated i.m., via the deltoid muscle. However, it has been recognized that a varying fraction of the vaccine injected i.m. can get into the blood in some individuals, sooner or later, via many possible routes.^26^

### Features of reactogenicity of Comirnaty in anti-PEG hyperimmune pigs

In sharp contrast to the study summarized above, where 5X HVD of Comirnaty caused no or minor physiological changes in naive pigs,^28^ in the present study using anti-PEG hyperimmune pigs, a 15-fold lower vaccine dose caused anaphylactic shock in 6 of 6 pigs with massive cardiopulmonary and other CARPA-specific physiological changes. This provides strong direct evidence that anti-PEG Abs, mainly anti-PEG IgM, play a causal role in vaccine-induced anaphylaxis, just as these Abs accelerate the mild HSRs to Doxil/Doxebo to anaphylaxis.^28^ There is, however, a major difference in the dose-dependence of Doxebo and Comirnaty-induced anaphylaxis. Namely, the PEG lipids in Doxil/Doxebo and Comirnaty are 2K-PEG-1,2-distearoylphosphatidylethanolamine (DSPE) and 2K-PEG-*N,N*-ditetradecylacetamide (ALC-0159), respectively, and the amount of 2K-PEG lipid administered to pigs with Doxil/Doxebo and Comirnaty were 25.0 and 0.68 µg/kg, respectively. This means that the amount of 2K-PEGylated lipid in Comirnaty was ∼37-fold less than that in equi-reactive Doxil/Doxebo. This points to increased sensitivity of the model for the vaccine-induced anaphylaxis compared to PEGylated liposomes, possibly because of the stronger C activation by Comirnaty compared to Doxil/Doxebo on a weight basis.

### Mechanism proposed

As mentioned earlier, mRNA-LNP vaccine-induced HSRs represent in most cases pseudoallergy,^9, 10, 15, 16, 19, 26^ proceeding without the involvement of specific IgE. The observations on significant in vitro C activation by Comirnaty in our previous,^33^ and in vivo C activation in the present study, wherein C3a rises in close parallelism with the clinical signs of anaphylactic reaction, provide direct evidence for the involvement of CARPA in the phenomenon. Just as described for Doxebo/Doxil-indued anaphylaxis,^28^ the chain reaction along the immuno-cardiovascular axis most likely involves a great number of redundant molecular and cellular interactions leading from C activation to variable vasoactivity in different organs (Fig 8).

**Figure 8.**
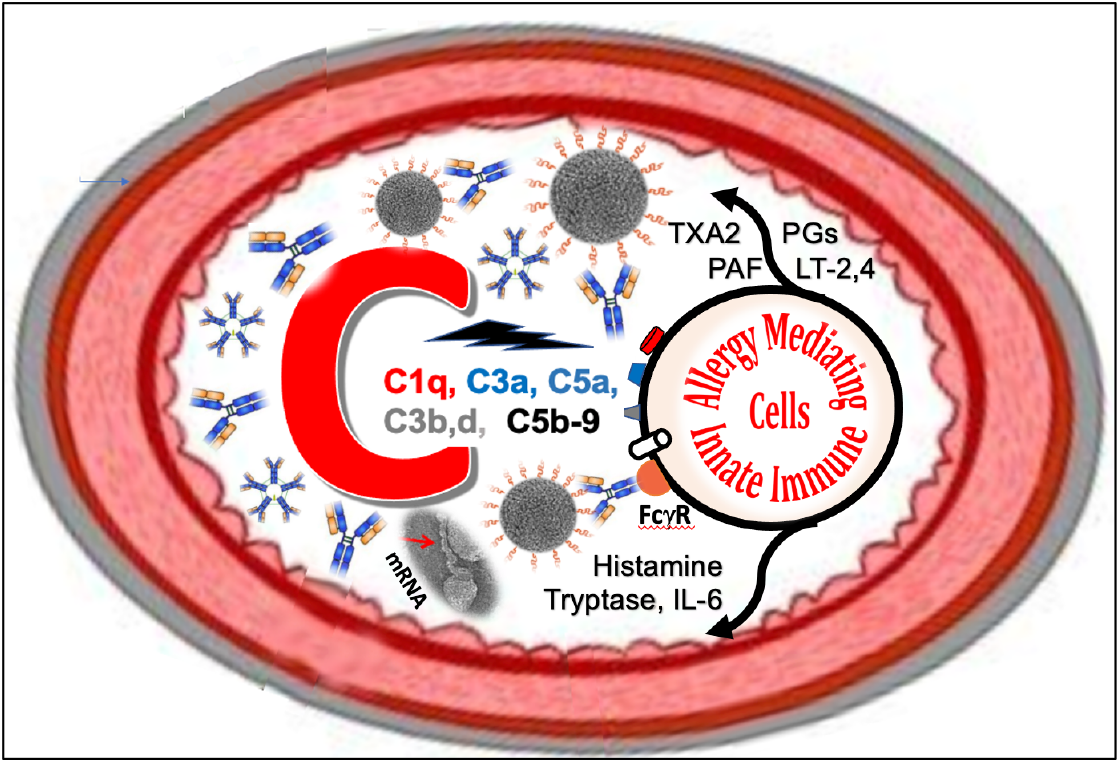
Possible mechanisms of HSRs/anaphylaxis by mRNA-LNP COVID-19 vaccines in pigs. After i.v. injection, the PEGylated vaccine NPs (solid black spheres with a crown) bind anti-PEG IgG and IgM Abs, which leads to C activation via the classical pathway. The liberated cleavage products C1q, C3a, C5a, C3b, -d, -dg and C5b-9 stimulate a variety of allergy-mediating innate immune cells (e.g., macrophages (PIM cells), mast cells, basophils, granulocytes, platelets) via different receptors (C1qR, C3aR, C5aR, CR1 (CD35), CR2 (CD21), CR3, CD11b/CD18, C5b-9R), illustrated with different colors and discussed in Ref. 33 in more detail. These signaling pathways represent CARPA, while the activations mediated by pattern recognition receptors, or pathogen-associated molecular patterns ((e.g., FcγRIIB/FcγR (CD32)/CD351), Toll-like receptor 2/6) represent C-independent pseudoallergy. The specified vasoactive secretory products, along with many more, explain the symptoms of HSR/anaphylaxis.^33^ The figure is a modification of Fig. 8 in Ref.^33^ with permission.

On the effector arm, we have evidence for the participation of the potent smooth muscle constrictor eicosanoid, TXA2, but the cell(s) releasing it without delay still remain(s) to be identified. TXA2 can be secreted by many cells including macrophages, endothelial cells, platelets and neutrophil granulocytes, but because pigs uniquely display pulmonary intravascular macrophages (PIM cells), their unique sensitivity to CARPA has been explained by these cells.^24, 34-38^ Likewise, we point to C activation as a major contributing mechanism to anaphylaxis, although the “double hit hypothesis”^46, 47^ predicts at least 2 simultaneous stimuli to be necessary to trigger an intense vasoactive mediator release by allergy mediating innate immune cells (Fig 8). Clearly, to achieve the goal of HSR-free vaccination, further exploration of both the afferent and efferent arms of vaccine-induced anaphylaxis will need to be performed, which is enabled by the present porcine model. To our best knowledge, it represents the first large animal model of drug- or vaccine-induced anaphylaxis.

## Supplement

**Figure S1.**
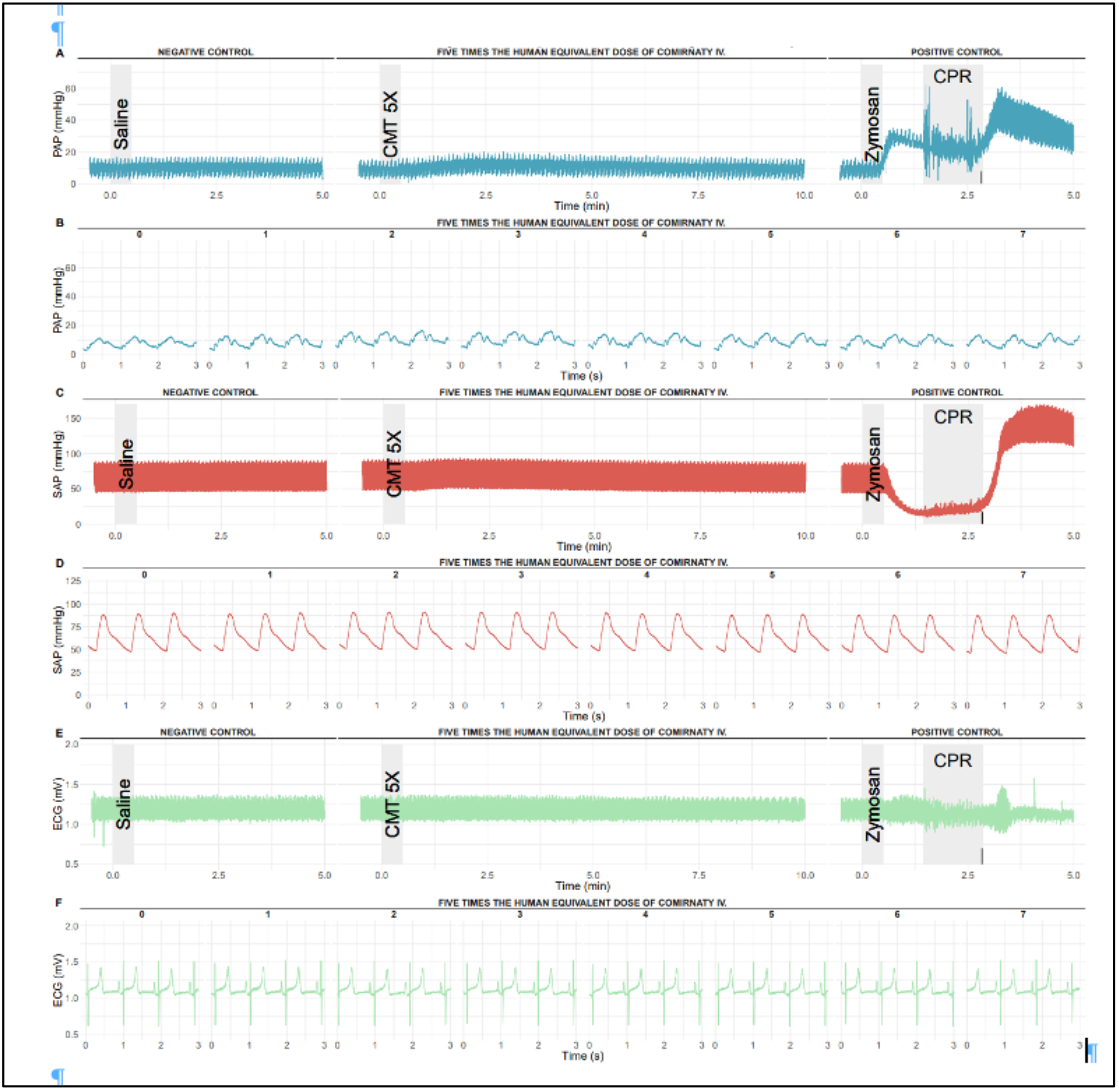
Typical real-time hemodynamic and ECG tracings in a pig injected i.v. with 5X HVD of Comirnaty after sham-immunization with PBS, as described in the Methods for Doxebo. Details of the Zymosan bolus and other abbreviations are the same as in Figures 2 and 3.

